# RNApysoforms: Fast rendering interactive visualization of RNA isoform structure and expression in Python

**DOI:** 10.1101/2024.11.06.622357

**Authors:** Bernardo Aguzzoli Heberle, Madeline L. Page, Emil K. Gustavsson, Mina Ryten, Mark T. W. Ebbert

## Abstract

**Motivation:** Alternative splicing generates multiple RNA isoforms from a single gene, enriching genetic diversity and impacting gene function. Effective visualization of these isoforms and their expression patterns is crucial but challenging due to limitations in existing tools. Traditional genome browsers lack programmability, while other tools offer limited customization, produce static plots, or cannot simultaneously display structures and expression levels. RNApysoforms was developed to address these gaps by providing a Python-based package that enables concurrent visualization of RNA isoform structures and expression data. Leveraging plotly and polars libraries, it offers an interactive, customizable, and faster-rendering framework suitable for web applications, enhancing the analysis and dissemination of RNA isoform research.

**Availability and implementation:** RNApysoforms is a Python package available at (https://github.com/UK-SBCoA-EbbertLab/RNApysoforms) via an open-source MIT license. It can be easily installed using the piip package installer for Python. Thorough documentation and usage vignettes are available at: https://rna-pysoforms.readthedocs.io/en/latest/.

## 1. Introduction

Alternative splicing is a fundamental biological mechanism that enhances genetic diversity by allowing a single gene to produce multiple RNA isoforms. This process, occurring in over 95% of human multi-exon genes^1,2^, results in mRNA variants that can have distinct and sometimes opposing functions. For instance, the *TRPM3* gene, which encodes human cation-selective channels, can be alternatively spliced to produce variants targeting different ions^3–5^. Similarly, *CASP3* has two transcript variants with opposing functions: one is pro-apoptotic, and the other is anti-apoptotic^6,7^. Alternative splicing is intricately regulated, resulting in a rich diversity of context-dependent RNA transcripts emerging from the same genetic code.

Traditional RNA sequencing methods that rely on short-read technologies face significant challenges in accurately reconstructing and quantifying these RNA isoforms. Short reads often do not span multiple splice junctions, making it difficult to assemble full-length transcript structures^8,9^. This limitation can lead researchers to aggregate all RNA isoforms into a single gene-level measurement, potentially obscuring important biological insights. The recent improvements and increased affordability of long-read sequencing platforms, such as PacBio and Oxford Nanopore, has revolutionized transcriptomics by enabling the sequencing of full-length transcripts. These technologies facilitate the discovery of novel isoforms and provide more accurate quantification of RNA isoform expression, leading to new insights in human genomics^10–12^, cancer^13–15^, Alzheimer’s disease^16^, and plant biology^17^.

Visualizing RNA isoform structures and expression patterns is crucial for interpreting transcriptomic data and forming hypotheses about gene function. However, many existing visualization tools lack flexibility or do not effectively compare transcript structures. Genome browsers like UCSC^18^ and IGV^19^ facilitate transcript visualization but are not easily customizable through programming. Other tools, such as IsoformSwitchAnalyzeR^20^, wiggleplotr^21^, and ggsashimi^22^, offer certain functionalities but provide limited options for adjusting plot aesthetics and comparing transcript structures. While SWAN^23^ offers customizable visualization within the Python environment, it has limitations in highlighting differences between isoforms and does not support interactive graphs.

Introduced in 2022, the ggtranscript^24^ R package provides a flexible and user-friendly framework for visualizing RNA isoform structures using ggplot2^25^, a widely adopted R-based data visualization library. As a ggplot2 extension, ggtranscript allows extensive customization and integrates well with other ggplot2 functionalities.

Despite its strengths, ggtranscript has notable limitations. It generates static plots without interactive capabilities, restricting users’ ability to dynamically explore RNA isoform structures—a significant drawback for web applications. Additionally, rendering complex plots can be time-consuming, making it less suitable for applications requiring rapid plot generation. Moreover, ggtranscript does not natively support concurrent visualization of RNA isoform expression levels alongside their structures, limiting its usefulness in studies focusing on differential isoform expression or usage.

To address these limitations, we developed RNApysoforms, a Python-based package designed to facilitate the simultaneous visualization of RNA isoform structures and expression levels. RNApysoforms aims to achieve three primary goals: (1) enable concurrent visualization of RNA isoform structures and their expression data; (2) provide a flexible framework for creating interactive plots, allowing users to explore RNA isoform structures and expression in variety of useful ways; and (3) enhance the dissemination of RNA isoform research findings through web applications by offering easy-to-use, interactive, customizable, and faster-rendering visualizations within the Python ecosystem. By leveraging the Python plotly^26^ and polars^27^ libraries, RNApysoforms improves performance and user experience in web-based environments, making it a valuable tool for researchers in genomics, transcriptomics, and related fields.

## 2. Implementation

RNApysoforms is a Python package that streamlines the analysis and visualization of RNA isoform data. It offers a powerful toolset for researchers by leveraging the polars library for efficient data manipulation and plotly for crafting interactive, customizable, and fast-rendering plots. The package provides functions to handle RNA isoform annotations—particularly ENSEMBL GTF files—and offers guidance for other transcript annotation formats. It supports processing count matrices and sample metadata files in various formats, such as .tsv/.txt, .xlsx, .csv, and .parquet. Utilizing polars-powered functions, users can easily wrangle transcript annotations, count matrices, and sample metadata to create customizable and interactive visualizations of transcript structures and expression data using plotly.

The read_ensembl_gtf() function reads an ENSEMBL^28^ GTF file and returns the data as a polars DataFrame, focusing specifically on ‘exon’ and ‘CDS’ feature types. It parses the ENSEMBL GTF file to extract key genomic features and attributes such as gene_id, gene_name, transcript_id, transcript_name, transcript_biotype, and exon_number. It performs several validation checks on the file and handles missing gene_name and transcript_name values by filling them with gene_id and transcript_id values where necessary.

The read_expression_matrix() function loads and processes an expression matrix, with options to merge it with metadata, perform Counts Per Million (CPM) normalization, and calculate relative transcript abundance. It reads an expression matrix file and, if provided, a metadata file, merging them on a specified sample identifier column. It supports CPM normalization and can also calculate relative transcript abundance based on gene counts when a gene identifier column is available. The resulting polars DataFrame is returned in long format, including expression measures, optional CPM values, relative abundances, and metadata.

The gene_filtering() function filters genomic annotations and optionally an expression matrix to focus on a specific target gene, with capabilities to order and select the top expressed transcripts. It narrows down the annotation DataFrame to entries matching the specified gene identifier. If an expression matrix is provided, it synchronizes the filtering to retain only transcripts present in the filtered annotation. Additionally, it offers the option to sort transcripts by their total expression levels and to keep a specified number of top expressed transcripts, streamlining the analysis of the most relevant transcripts associated with the target gene.

The calculate_exon_number() function assigns exon numbers to exons, coding sequences (CDS), and introns within a genomic annotation dataset based on transcript structure and strand orientation. It processes a genomic annotation DataFrame, numbering exons sequentially within each transcript while accounting for strand direction—incrementing from the 5′ to 3′ end on the positive strand and decrementing on the negative strand. CDS and introns receive exon numbers based on their overlap with or adjacency to the numbered exons. This function facilitates support for annotation file formats that may not contain exon numbers.

The to_intron() function converts exon coordinates into corresponding intron coordinates within a genomic annotation DataFrame. It identifies introns by calculating the genomic intervals between consecutive exons for each transcript, determining the regions between exons that constitute introns. The function returns a DataFrame containing both exon and intron coordinates, facilitating proper intron visualization.

The shorten_gaps() function reduces intron and transcript start gaps between exons in genomic annotations to a specified target size, enhancing the visualization of exons and/or CDS regions. It improves the clarity of transcript visualizations by minimizing the visual space occupied by long introns and aligning transcripts for consistent rescaling, all while maintaining the relative structure of the transcripts. This function facilitates the creation of more interpretable and visually appealing representations of transcript structures.

The make_traces() function generates plotly traces for visualizing transcript structures and/or expression data, enabling detailed and customizable genomic data visualization. It processes genomic annotation and/or expression data to create plotly traces suitable for plotting transcript structures alongside expression metrics. It supports extensive customization of plot aesthetics—including colors, line widths, plot styles, and annotations—allowing users to tailor the visual representation to their specific needs. By returning a list of traces that can be directly used in plotly figures, it facilitates the creation of comprehensive visualizations of genomic features such as exons, introns, and coding sequences (CDS), as well as associated expression data.

The make_plot() function creates a plotly figure that can feature either single or multiple panels, designed to visualize transcript structures and/or expression data comprehensively. It builds a customizable and interactive figure with horizontally arranged subplots, accepting a list of plotly traces that represent transcript structures and expression metrics. These traces are organized into subplots, with the layout, axes, and overall appearance tailored to user-defined parameters. This flexibility enables detailed and adaptable visualizations, enhancing the interpretation of complex transcriptomic data.

RNApysoform functions exclusively support polars DataFrames. However, pandas and pyarrow are included as package dependencies, allowing users to manipulate their data using the more popular—but slower— pandas library. Users can then convert their pandas DataFrames to polars DataFrames before utilizing the RNApysoform functions. Additionally, the package includes kaleido as a dependency, enabling users to save their generated plots as static images—a feature not natively supported by plotly, which typically allows saving plots only in interactive formats like HTML.

## 3. Execution time comparison

We evaluated the performance of RNApysoforms compared to ggtranscript by measuring the time required to render and save 1,499 plots covering all genes and transcripts from human chromosomes 21 and Y using the ENSEMBL GTF release 107. Each software package was tested over 100 iterations (n=100), with each iteration generating the complete set of plots. RNApysoforms saved plots in HTML format (natively supported by plotly), while ggtranscript used PDF format (natively supported by ggplot2). Due to differences in format and interactivity, RNApysoforms plots were significantly larger in size, occupying 6.5 GB for 1,499 plots compared to ggtranscript’s 94 MB. Though, because RNApysoforms is typically rendered directly on the webpage, it does not generally need to generate output files. The execution time comparison scripts were run sequentially on the same compute node at the University of Kentucky’s Morgan Compute Cluster, utilizing 1 CPU and 4 GB of RAM. All data, code, outputs, and the software container required for this analysis are accessible via the GitHub repository here (https://github.com/UK-SBCoA-EbbertLab/article_analysis_RNApysoforms).

RNApysoforms demonstrated a median runtime of 121.25 seconds (interquartile range [IQR]: 119.69–122.71 seconds), significantly outperforming ggtranscript’s median runtime of 206.87 seconds (IQR: 185.34–226.44 seconds). This substantial difference translates to RNApysoforms being approximately 70% faster, with a median time reduction of 85.63 seconds (**Figure 1b**). Statistical analyses revealed significant deviations from normality in the runtime data for both tools, as indicated by Shapiro-Wilk tests (RNApysoforms: W=0.826, p=1.67×10^−9^; ggtranscript: W=0.953, p=0.0014), and unequal variances confirmed by Levene’s test (F=238.66, p=7.55×10^−36^). Consequently, we employed a non-parametric permutation test on the median difference, which, using 10 million resamples, revealed a significant difference in execution times favoring RNApysoforms (p=2×10^−7^). Although a smaller p-value might be obtained with more resamples, increasing the number of resamples was not practical due to the considerable computational resources required. To further quantify this advantage, we built a 95% bootstrap confidence for the difference in execution time using 10 million resamples and the interval was [-94.59, -77.05] seconds, underscoring RNApysoforms’ substantial performance edge.Furthermore, an effect size of -1.0 (Cliff’s delta) indicates a large and consistent performance difference. Collectively, these results establish RNApysoforms as a significantly faster tool for visualizing RNA isoform structures in large-scale genomic datasets.

**Figure 1:**
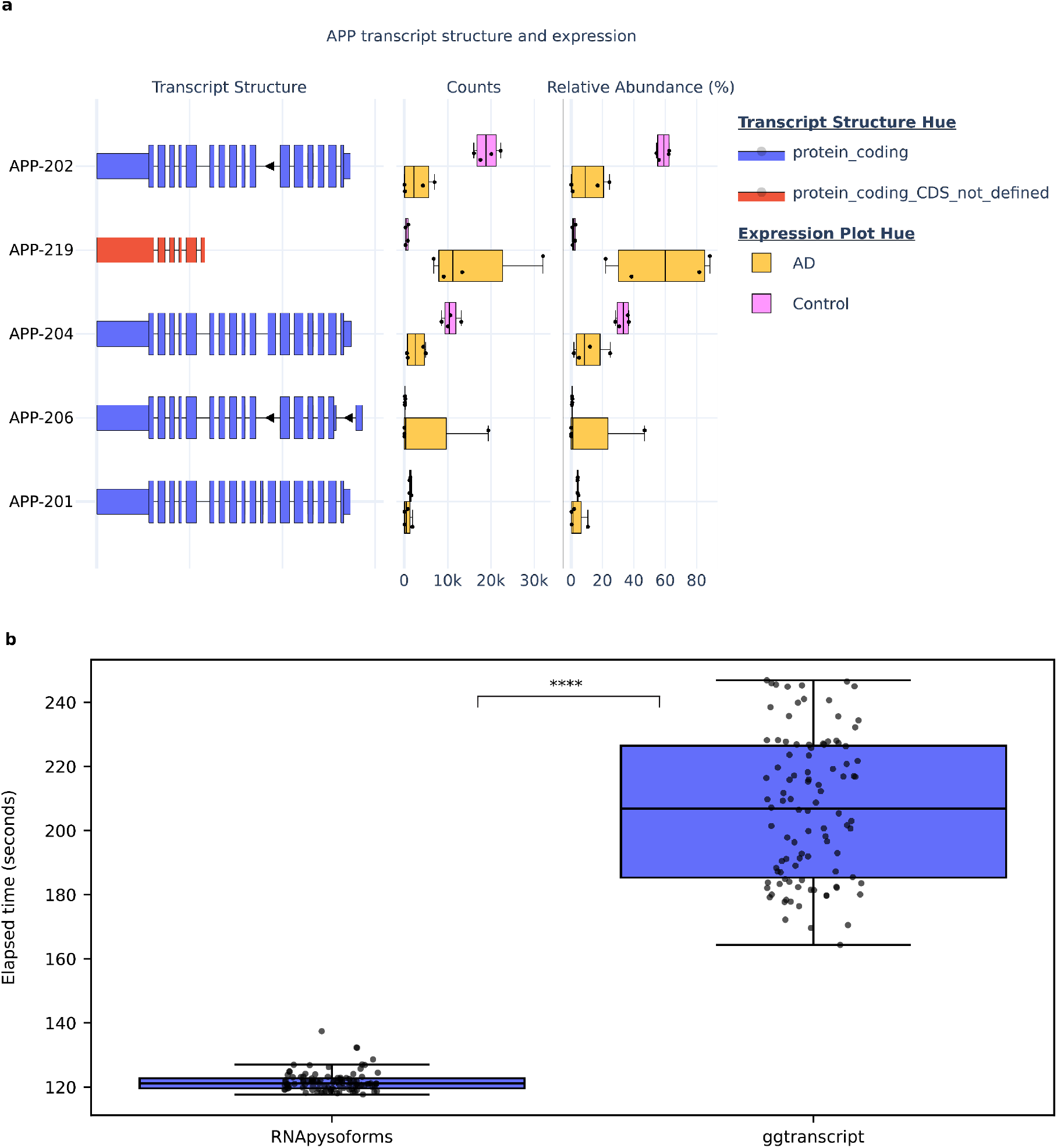
RNApysoforms visualization example and execution time comparison with ggtranscript. **a**, Example plot generated by RNApysoforms illustrating the structure and expression of five APP RNA isoforms in a test dataset comparing Alzheimer’s disease (AD, n=4) with control (n=4) samples. An interactive version of the plot is available at the bottom of [our vignette](https://rna-pysoforms.readthedocs.io/en/latest/examples/14.using_transcript_name_instead_of_transcript_id.html). **b**, Boxplot showing the time taken by RNApysoforms and ggtranscript to render 1499 plots, encompassing all genes and transcripts from human chromosomes 21 and Y in ENSEMBL GTF release 107. Each package ran 100 iterations (n=100) with 1499 plots generated per loop. RNApysoforms achieved a median completion time of 121.25 seconds, 70% faster than ggtranscript’s median of 206.87 seconds. A permutation test on the median execution time difference yielded a significant p-value = 2×10^−7^, with a 95% bootstrap confidence interval for the execution time difference of [-94.59, - 77.05] seconds. All boxplots in this panel follow the following format: median (center line), quartiles (box limits), 1.5 × interquartile range (whiskers). **** p-value < 0.0001.

## 4. Conclusion

RNApysoforms successfully addresses the limitations of existing RNA isoform visualization tools by providing a Python-based package that enables simultaneous visualization of RNA isoform structures and expression levels. By leveraging efficient data manipulation with polars and interactive plotting with plotly, it offers a flexible framework for creating customizable and interactive plots. Performance evaluations demonstrate that RNApysoforms significantly outperforms ggtranscript in rendering and saving plots, making it a faster and more efficient tool for large-scale genomic datasets and web applications. This enhanced performance, combined with its ease of use and integration into web applications, makes RNApysoforms a valuable resource for transcriptomics research.

## Acknowledgments

We appreciate the contributions of the Sanders-Brown Center on Aging at the University of Kentucky. We would like to thank the University of Kentucky Center for Computational Sciences and Information Technology Services Research Computing for their support and use of the Morgan Compute Cluster and associated research computing resources. We would like to thank Singularity Sylabs for providing support and extra cloud storage for our software containers. This research was funded in part by Aligning Science Across Parkinson’s [grant numbers: ASAP-000478 (M.R.), ASAP-000509 (E.K.G. and M.R.) through the Michael J. Fox Foundation for Parkinson’s Research (MJFF).

